# A microfluidic droplet array demonstrating high-throughput screening in individual lipid-producing microalgae

**DOI:** 10.1101/2022.05.05.490790

**Authors:** Guoxia Zheng, Furong Gu, Yutong Cui, Ling Lu, Xuejun Hu, Lin Wang, Yunhua Wang

**Affiliations:** Environmental and Chemical Engineering Institute, Dalian University, Dalian, 116622, China; Dalian Key Laboratory of Oligosaccharide Recombination and Recombinant Protein Modificatio n,Dalian, 116622, China; Medical School, Dalian University, Dalian, 116622, China

## Abstract

Microalgae are a group of photoautotrophic microorganisms which could use carbon dioxide for autosynthesis. They have been envisioned as one of the most prospective feedstock for renewable oil. However, great endeavors will still be needed to increase their economic feasibility; the screening of competitive species and suitable culture conditions are such issues. To greatly accelerate these rather laborious steps and also improve their experimental lump-sum-manner, we developed a microfluidic droplet-based 2×10^3^ resolution “identification card”, which allowed high throughput real-time monitoring of individual algae among population. A novel fluid-blocking-based droplet generating and trapping performance were integrated in the platform which made it excellent in operational simplicity, rapidity and stability and full of the potentials in single-cell-isolation/screening. The developed platform was successfully used to screen three unicellular algae, namely, *Isochrysis zhanjiangensis, Platymonas subcordiformis* and *Platymonas helgolandica var. tsingtaoensis*. In situ bioassays of the lipid accumulation and cell proliferation at single cell level for interspecies comparison were possible. Nitrogen stress condition can be indentified that induce positive-skewed frequency distribution of lipid content.

## 1. Introduction

In past decades, due to the continuous rising of crude oil prices and global warming by green house effect, development of alternative substantial energy has attracted more and more attention [1, 2]. Biofuels, including biodiesel production from microalgae, have been increasingly considered potential substitutes for fossil energy for their renewable feature [3, 4]. However, the high cost of biofuels limits their availability of commercialized production [5]. It is urgently needed to screen competitive strains (for example, lipid-rich and fast-growing strains), and to improve large-scale cultivation based on understanding the effects of various culture conditions, such as light intensity, nutrition, carbon dioxide, etc [6, 7]. Usually, these works are accomplished by using traditional flask-type algae culturing and lipid analyzing technique [8, 9]. These methods are effective. However, traditional flask-based bioreactors are not qualified for high-throughput screening. To improve biodiesel yields, combination of various culturing factors and numerous microalgae strains, both natural and engineered, are needed to be evaluated. The workload goes far beyond hundreds of times throughput capacity of the currently available culturing systems. Obviously, an integrated biological/biochemical platform that can quickly screen to identify a single microalgal cells and culture conditions for high lipid production and fast growth, while in a high-throughput manner, could significantly accelerate the process of industrialization of algae derived biofuel.

To date, a large number of delicate implements based on advancing microfabrication technology have emerged for single-cell analysis [10]. For example, microwell arrays enable cells to be isolated and observed in individuals [11]. However, this method has not been facilitated algae screening, which basically attribute to the minimum cell size and floating, even motile nature of microalgae. Optical tweezers [12], magnetic tweezers [13] and magnetic tweezers [14] could achieve extremely high resolution and 3D manipulation, but they rely heavily on sophisticated system and high energy/power. Microfluidic chip devices which have received great interests recently for their capability to precisely manipulate fluids at picoliter, nanoliter or microliter level as well as to integrate multiple analysis steps within micrometer-scale microstructures could be an ideal alternative [15, 16]. Especially, droplet-based microfluidics has dimensional scaling benefits and enables precise generation and repeatable operations of fluids into droplets [17]. Aqueous droplets could be generated by water-in-oil emulsion process and suspended in continuous oil phase. Each droplet, with volume of pico to nano-liter scale, could encapsulate one or more cells and be used as an independent bioreactor, allowing for massive parallel processing [18-21]. It has been successfully applied in various biological/biochemical screening applications (eg. drug candidate screening, bioactive molecular synthesis and enzyme investigation, *etc*.) [22-26]. However, a microfluidics droplet-based platform with all algae screening steps (i.e. droplet generating and cell encapsulating, droplet trapping and immobilization, on-chip cell growth, staining and in-situ bioassays) occurring seamlessly on-chip in a high-throughput manner, whether from chip manufacture or fluid manipulation, is difficult to achieve [27, 28]. Thus, despite those tremendous potentials, the droplet-based algae screening platform has not been used widely.

In this research, we provided a microfluidic droplet-based high-throughput screening platform for real-time analyzing of algal lipid accumulation and growth at individual cells. Different from previous works, a novel fluid-blocking-based droplets generating and trapping method, combining with diffusion-based on-chip BODIPY staining, was developed in this study, which made the chip excellent in operational simplicity, rapidity and stability. Thousands of nanoliter-sized droplets generated and trapped with high uniformity, could sever as an “identification card” which contains several biometric signatures at single cell level, such as metabolism (i.e. lipid accumulation), phenotypes and proliferation, *etc*. In-situ observation of cellular lipid accumulation and proliferation in single microalgal cells within droplets for interspecies comparison could be achieved.

As shown in Fig. 1, the “identification card” is made up of two layers: an upper stratum functionalis and a substrate (Fig.1a). Upper stratum functionalis consists of two functional units: a fluidic-blocking network and a trap array. The fluidic-blocking network includes a series of parallel fluidic-blocking channels (100μm wide and 100μm high) in which the oil phase flows to block aqueous phase into droplets. The trap array consists of 2112 round-shape trap chambers for droplet trapping and immobilization. These chambers were linked in series by algae transportation channels (50μm wide and 50μm high). . Each trap chamber (240μm of diameter and 50μm high) is centered between two parallel fluidic-blocking channels and were connected by stop-flow junctions where the two-phase interface exists. The height of the trap chambers and flow-stop junctions was lower than that of the fluidic-blocking channels.

**Fig. 1.**
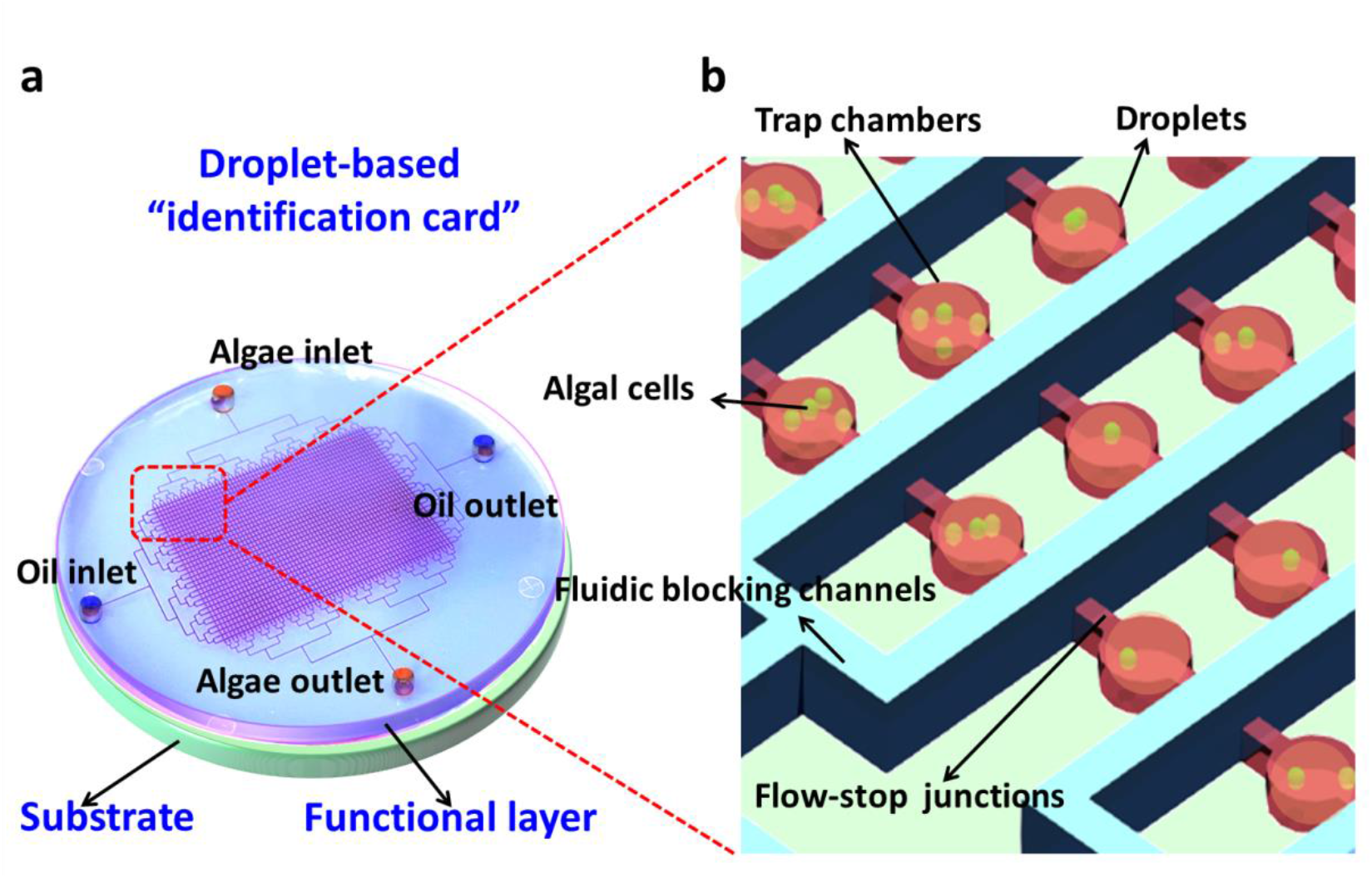
Schema of the droplet-based microfluidic platform for high-throughput screening in individual microalgae; (a) Structure of the droplet array chip, which consists of two layers: an upper stratum functionalis and a substrate; These two layers are enclosed together to form ―identification card”. (b) The magnified view of functional layer, which consists of a fluidic blocking network (blue) and a trap array (red); Each round trap chamber is centered between two parallel fluidic blocking channels and connected them by stop-flow junctions where the two-phase’s interface exists.

Both the functional layer and the substrate layer were fabricated in polydimethylsiloxane (PDMS, Sylgard 184, Dow Corning, USA) as described previously [29]. To operate the device, the syringe pumps (Harvard, USA) were used to actuate the dispersed phase (aqueous solution with f/2 medium) or continuous phase (fluorinated oil FC-40) to flow through the microfluidic chip. The operation procedures were described as follows: (i) Chip preparation. The syringe pumps were connected with two inlet holes of the chip by two PEEK tubes, respectively; (ii) The oil outlet of fluidic-blocking network was sealed and the algae outlet of the trap array was left open. Then the water solution was injected in the chip to fill the trap array only. (iii) Open the oil outlet of fluidic blocking network and seal the algae outlet of trap array, and then continuous phase was injected in the chip to fill the fluidic-blocking network. (vi) The middle located dispersed phases were sheared into droplets by their two lateral continuous phases at stop-flow junctions. And then, the generated droplets were trapped and immobilized in each of trap chambers. Meanwhile, algal cells were encapsulated in the droplets.

The capability of the device to encapsulate single algae cell into droplet was investigated. In this assay, an algal cells suspension in f/2 medium was used as dispersed phase and was injected at a flow rate of 450 μl min^-1^.The density was adjusted to7-8 cells μl ^-1^ to ensure that a single algal cell be encapsulated into one droplet. As illustrated in figure 2, uniform aqueous droplets were trapped in an array format (Fig 2a), encapsulating of single algal cells (Fig 2b). The developed platform was capable of encapsulating individual algal cells into array droplets with 7–10% probability and generally no multi-cell-containing droplets. The confining ability of the droplet-based platform was investigated by encapsulating motile algal cell (*P*.*helgolandica*) into droplet. In such issues, droplets functioned as “mini-cages” that confine “running” cell inside. It enabled not only an isolation of each “running” sample and prevention of cross-contamination, but also a more accurate quantitative microscopic inspection than that done on plates. And thus, the developed droplet-based platform is also especially beneficial when performing cell screening with motile samples. Noticeably, the current fluidic-blocking-based droplet generating and trapping method is extremely simple, rapid and stable as compared with most of traditional channel-based droplet manipulation methods, such as “T” or “Y” junction droplet generating, floatage-based or resistance-based droplet trapping, *etc* [27, 29]. Additionally, the trap array are isolated with each other at the top layer, which makes it simple to accomplish the biometric signatures of numerous individual algal cells by a CCD camera without changing the location of the chip. It also allowed for integrating an extra picking operation by use of syringe extraction for separating the desired algal species from its trap chamber without disturbing the neighbors. And thus, the established droplet-based platform, serving as a smart “identification card”, enabled a high throughput real-time monitoring of individual algae among population, allowing multiple algal screening, bioassays and isolation at the same time.

**Fig. 2.**
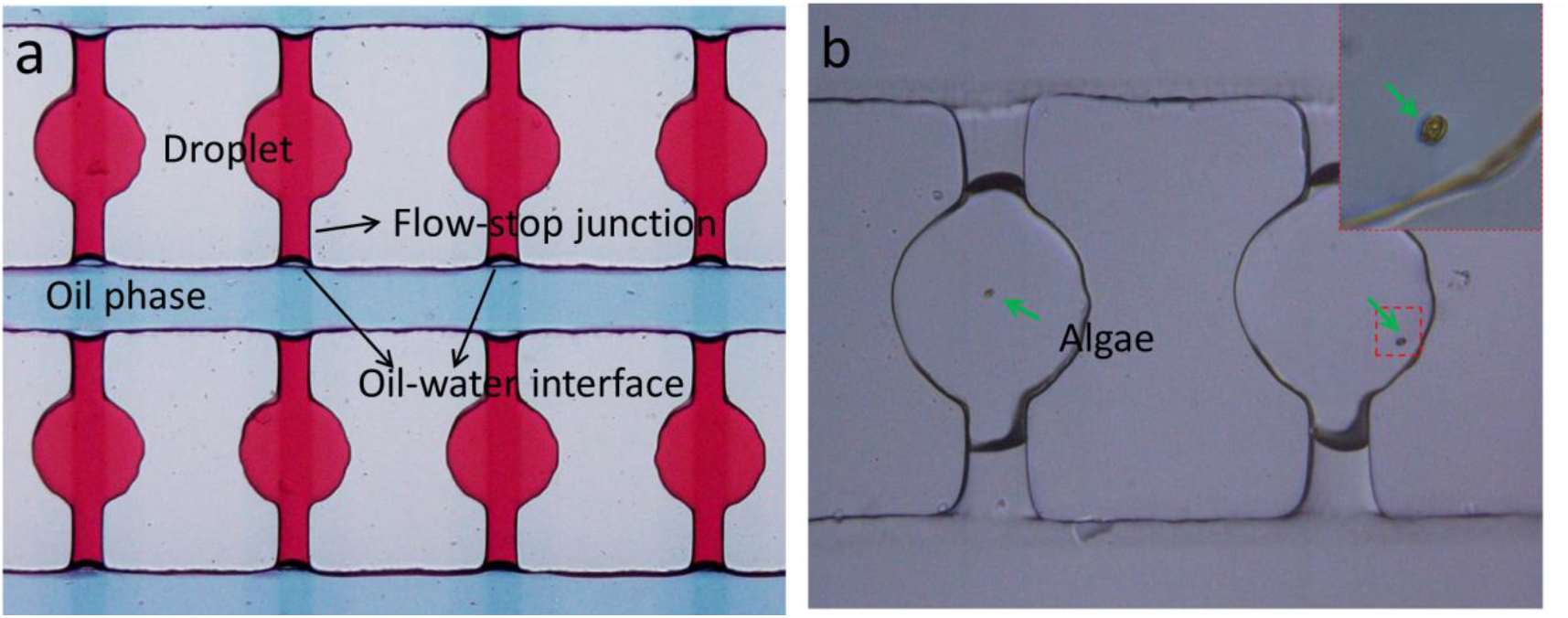
The fluidic-blocking-based microfluidic platform for trapping of droplet array. (a) Photographs of the microfluidics trapping of droplets;(b) A microfluidics trapping of droplets encapsulated with single microalga. The inset is the encapsulated single alga (magnified view).

Traditionally, the algal lipids were determined gravimetrically, which required at least 10-15mg wet cells [35]. These results represented the average lipid production level of a population and the heterogeneity among individual cells could not be identified, while the fluorescence measurement of neutral lipid displaying lipid content of every single cell will be more suitable for single-cell screening [36]. Especially, emission spectrum of BODIPY staining (green) is separated from that of chlorophyll autofluorescence (red), which allows effective cell phenotype discrimination [37]. Additionally, BODIPY is a non-destructive staining reagent, thus further analysis on stained cell are also possible [38]. However, on-chip fluorescent staining is not easily 100% achieved in droplet-based device that need to synchronize a cell-containing droplet with a fluorescence dye-contain droplet to obtain one-to-one merging by an electric pulse [27]. And therefore, a simple on-chip staining method was applied in this study. It was achieved by diffusing BODIPY from the oil phase to the trapped aqueous droplets. Figure 3 depicted the diffusion of BODIPY from the oil phase to the trapped aqueous droplets. The fluorescent intensity changes at 0, 5min, 15min and 60min were analyzed, which were calculated for the pixels (distance) from one of the fluid-blocking channel to the trap chamber. From 15 min time point, the intensity profile across the chamber maintained as nearly 3/7 of that across the fluidic-blocking channel. It implied that the BODIPY distribution ratio in oil phase was at least 2.3 times as much as that in liquid phase, which should attribute to the lipophilic character of the dye [37].

**Fig.3.**
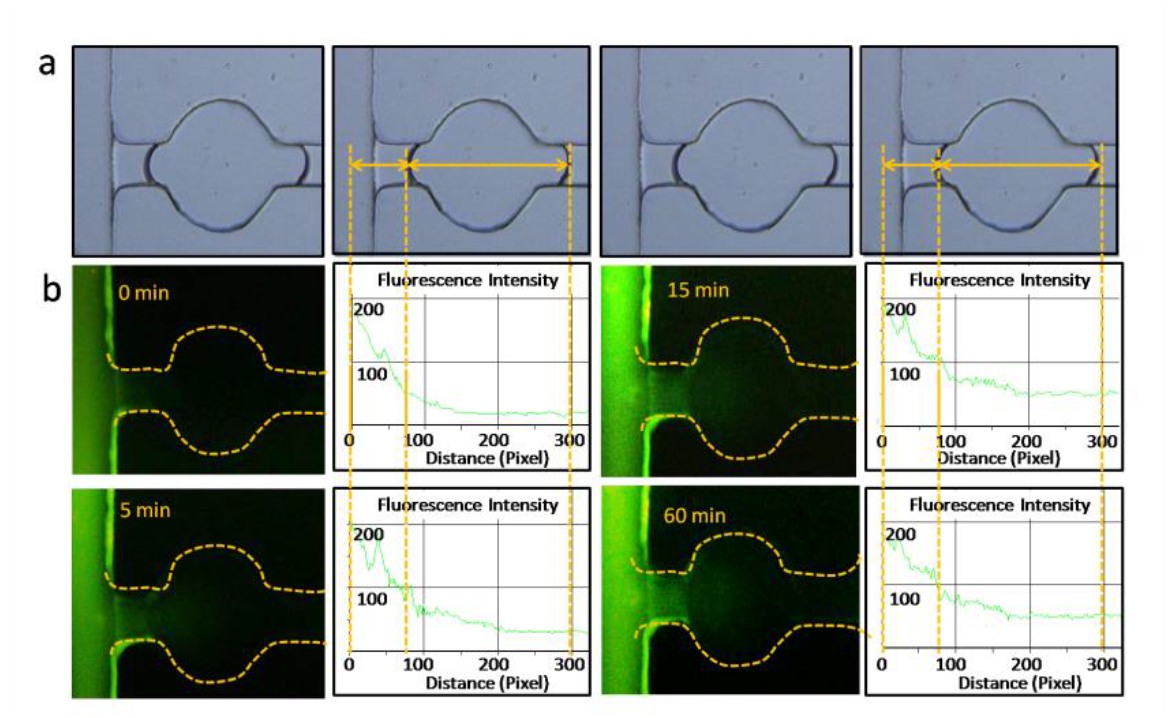
Characterization of the diffusion of BODIPY from the oil phase to the trapped aqueous droplets. (a) Images were processed to obtain the changes in fluorescent intensity at 0, 5min, 15 and 60min time point, respectively. Fluorescent intensities were calculated for all pixels (distance) across the length from one of the fluid-blocking channel to the trap chamber using Image-Pro Plus software.

To further characterize on-chip BODIPY-staining capability, N-deplete *P*.*helgolandica* cells (0% nitrogen) were encapsulated in single cell level within one droplet. *P*.*helgolandica* were tested here for their relatively large cell size and rigid cell wall which make them well suited for determining the maximum dye quality. Algal cells were encapsulated in droplets. Various concentrations of BODIPY 493/503 were diffused from oil phase to the trapped aqueous droplets, respectively. After 30 min incubation in the dark, imaging experiments were conducted for the single-cell-containing droplets by use of Olympus IX-71 fluorescence microscope (Olympus, Japan) equipped with DP73 CCD camera. Three different BODIPY concentrations (1µg ml^-1^, 1.5 µg ml^-1^ and 2.5µg ml^-1^ in fluorinated oil FC-40)were tested. Figure 4a shows the representative photographs of individual stained *P*.*helgolandica*. Their cellular lipid bodies, almost entirely composed of nonpolar triacylglycerols (TAGs) [39], enabled an easy visualization as light green by staining with a lipophilic BODIPY, even when merged with red chlorophyll fluorescence. As shown in figure 4b, the variations in fluorescence intensities could be observed depending on various BODIPY concentrations. When using 1.5 µg ml^-1^ BODIPY, general similarity of staining was achieved with 2.5 µg ml^-1^ staining. Considering that 2.5 µg ml^-1^ staining is excessive and back-diffusion performance would be needed to eliminate its disturbance in visual detection. And thus, 1.5 µg ml^-1^ staining was applied as maximum dye concentration for all algae strains.

**Fig.4.**
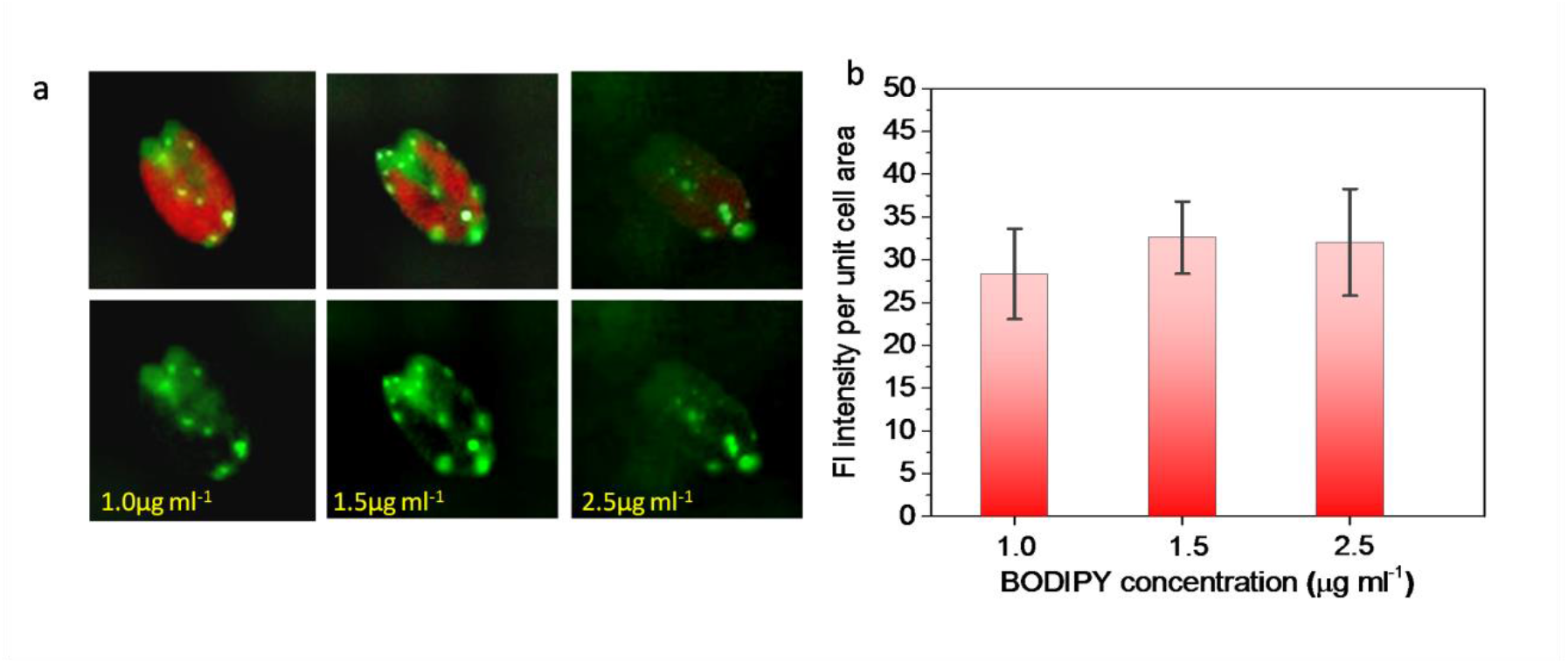
Characterization of BODIPY-staining of algae within droplet. (a) BODIPY stained oil bodies (light green) in *P*.*helgolandica* and their merged images with chlorophyll autofluorescence (red); (b) Analysis of average values of fluorescence intensity of lipid bodies stained with different concentrations of BODIPY.

Lipid productivity is determined by the growth rate and lipid production of microalgae [5]. Identification and isolation of strains with such high performance is critical but challenging, especially in a high-throughput manner [7, 9]. In traditional library screening, cell populations are diluted and plated on media plates. Cells which exhibited the desired properties (e.g., lipid-rich and yet fast-growing cells) are then isolated, proliferated, and identified for germplasm certification [8]. Although useful, this method is very labor-intensive and low throughput. And therefore, we have endeavored to develop a high-throughput microfluidics droplet-based microalgae screening platform that is capable of forming a ∼2×103 “identification card” with single-cell resolution within several seconds.

To demonstrate on-chip library screening functionality in oleaginous microalgae, cell populations of three species of *I. zhanjiangensis, P. subcordiformis* and *P. helgolandica var. tsingtaoensis* were tested in the developed platform. Droplets containing single cell were generated on-chip. After staining with BODIPY, single-cell-containing droplets were measured in situ for lipid contents. Images of green fluorescence in selected single cell of each species were captured. Figure 5a demonstrated the average fluorescence intensities per unit cell area in the three microalgal species within droplets, which is indicative of their average lipid contents. As it illustrated, intracellular lipid levels were distinguishable, although cells were encapsulated in the micro-droplet. *I. zhanjiangensis* exhibited higher lipid content than other two and can be the selection for biodiesel production. For further validation of on-chip screening data, off-chip oil production measurements were performed by total lipid extraction. As expected, the same conclusion was gained from off-chip experiments that *I. zhanjiangensis* exhibited more desired lipid-rich property than the other two (Fig. 5b). It was suggested that an inter-specific difference of lipid accumulation among oleaginous microalgae could be clearly indentified using fluorescence-activated cell sorting within the developed droplet array platform.

**Fig.5.**
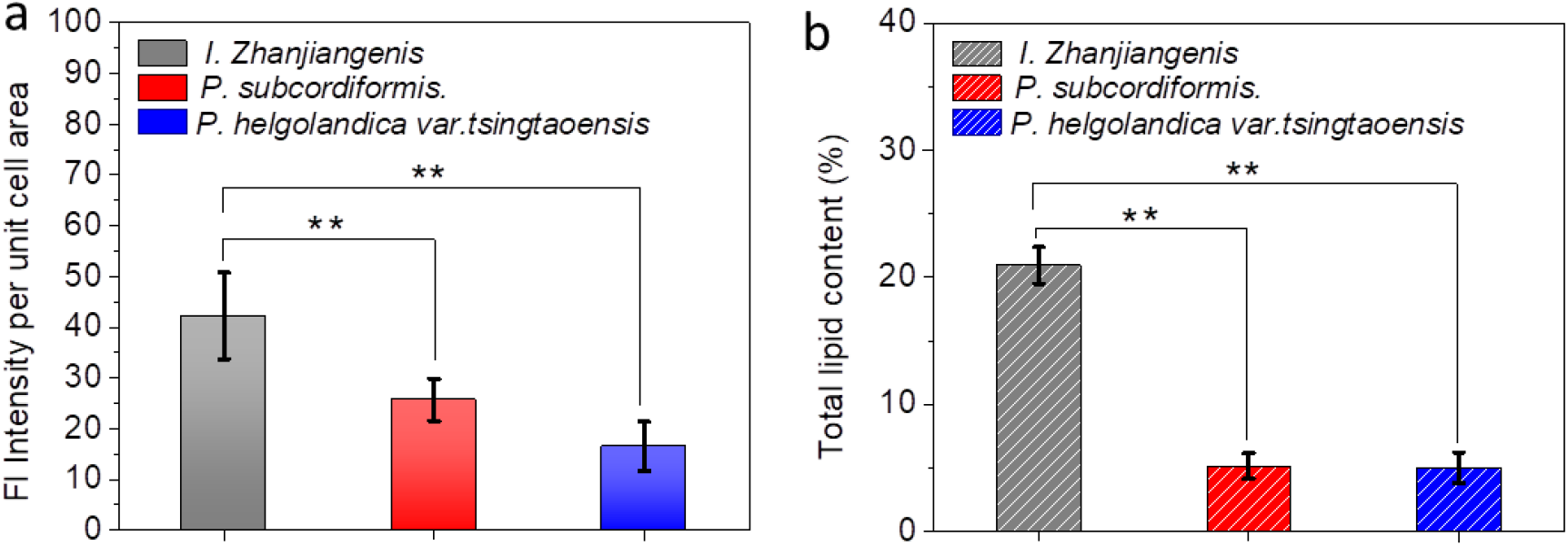
On-chip and off-chip analyzing of lipid content of different algae species. (a) The average fluorescence intensity profiles of each species of the three microalgae within droplets (*n* = 12). (b) Off-chip measurements of oil accumulation in each species of the three microalgae using gravimetrical method.

In order to validate the capability of the platform in quantifying the algal proliferation and also the on-chip cell viability, three tested species were grown for 7 days in the droplet-based screening platform. As we would expected, all these three species have successfully proliferated from single cell to single colony. Their colony sizes were calculated by counting the cell numbers within droplets using image J software. The final population size of the three algae, namely *I. zhanjiangensis, P. subcordiformis* and *P. helgolandica var. tsingtaoensis* declined in turn. The same trend was also found in off-chip experiments. From these growth data, *I. zhanjiangensis* was the faster-growing strain which should be the most suitable for large scale culture for biodiesel production among the three strains.

## 4. Conclusions

We proposed a novel high-throughput microfluidic droplet-based screening platform for analyzing interspecific difference and intraspecific inhomogeneity in individual lipid-producing microalgae. By comprehensive utilization of the temporary fluidic valve, density variations and also surface-tension equilibrium between oil and liquid phase, ∼2×10^3^ droplets were successfully generated within several seconds and almost simultaneously trapped in high uniformity during the process of the oil phase FC-40 blocking the aqueous phase. This so-called fluidic-blocking-based droplet manipulation platform is quite simple, stable and easy to operate as compared with most of traditional microludic droplet-based device. It enabled the encapsulation of single algal cell into droplets with 7–10% probability and generally no multi-cell-containing droplets. This single cell encapsulation combined with diffusion-based on-chip staining has facilitated real-time analyzing of multiple individual algal cells in a droplet-array-format, allowing high-throughput algal screening.

Similar to off-chip experimental data, the lipid fluorescence intensity and algae growth varied among the three tested species, *I. zhanjiangensis, P. subcordiformis* and *P. helgolandica var. tsingtaoensis*. The proposed high-throughput microfluidic droplet-based platform is now ready for use in algae screening for development of renewable energy and represents a multipurpose tool for bioengineering research at single cell resolution.

## Acknowledgements

This work was supported by National Natural Science Foundation of China (No. 41476085 and No. 81471807) and General program of Liaoning Science and Technology project (No. 2021-MS-345).

